# The neuropeptide alpha calcitonin gene-related peptide impairs load-stimulated proteoglycan production of human chondrocytes via WNT activity

**DOI:** 10.1101/2024.10.11.617807

**Authors:** H.F. Dietmar, N. Hecht, C. Binder, S. Schmidt, T. Walker, S. Grässel, W. Richter, S. Diederichs

**Author notes:** Corresponding author: PD Dr. Solvig Diederichs, Research Centre for Molecular and Regenerative Orthopaedics, Department of Orthopaedics, Medical Faculty Heidelberg, Heidelberg University, Schlierbacher Landstr. 200a, 69118 Heidelberg, Germany.

## Abstract

**Objective:** Novel targets for osteoarthritis therapy are urgently needed, and sensory nerve fibres and their neuropeptides are increasingly recognised for their contribution to structural aspects of joint pathology. The nociceptive sensory neuropeptide alpha calcitonin gene-related peptide (αCGRP) was previously detected in synovial fluid and serum of osteoarthritis patients and was also described as trophic factor for chondrocytes, affecting ECM organisation and biomechanical properties. Here, we investigated the potential of αCGRP to alter the chondrocyte mechanoresponse, and thus to affect the resilience of cartilage towards mechanical loading.

**Methods:** Tissue-engineered neocartilage based on human articular chondrocytes was treated with 1µM αCGRP for 24 hours and subjected to an anabolic loading protocol (intermittent dynamic compression, 1Hz, 25%) for the last 3 hours before analysing its molecular mechano-response.

**Results:** Mechanotransduction was largely unaltered by αCGRP as demonstrated by ERK activation and stimulation of mechano-regulated gene expression, yet load-stimulated glycosaminoglycan synthesis was disturbed by αCGRP. Presence of αCGRP did not affect stimulation of *WNT5A* expression by loading, but decreased *DKK3* expression under loading. Importantly, WNT inhibition prevented the negative effect of αCGRP on load-stimulated glycosaminoglycan synthesis.

**Conclusion:** Identifying *WNT5A* as a novel mechano-response gene, we reveal that αCGRP can block the load-stimulated proteoglycan production of human chondrocytes in the presence of WNT pathway activity. Thus, our data propose a novel, negative role for the pain-mediator αCGRP in the cartilage loading response, compromising its resilience to loading. Overall, our study implicates αCGRP as a potential target for osteoarthritis treatment, but also in patient stratification.

## INTRODUCTION

Osteoarthritis (OA) is a leading cause of disability, with 7.6 % of the global population (nearly 600 million people) living with some form of OA in 2020^1^. During OA, the structural and functional properties of joint tissues, including articular cartilage, are progressively compromised, resulting in symptoms such as joint stiffness, immobility and pain. Whilst the burden of disease is predicted to rise further by 2050, there is no effective cure for OA and despite intense efforts, no disease-modifying osteoarthritis drugs (DMOADs) have been approved by regulatory agencies to date^2^. Thus, there is an urgent need to identify novel potential targets for OA prevention and therapy. Importantly, physiological mechanical stimulation contributes to cartilage homeostasis and maintenance^3-5^; however, cartilage degeneration during OA impairs this functional capacity to withstand and adequately transmit mechanical forces during joint movement, affecting its resilience towards mechanical loading^2^. An improved understanding of the molecular chondrocyte loading response and how it is affected by degenerative changes within the joint could therefore reveal novel approaches to enhance the resilience of (OA) cartilage towards mechanical loading and thereby slow or prevent disease progression.

Sensory nociceptive nerve fibres are increasingly recognised for their contribution to joint pathology, but whether they compromise the mechanoresponse of cartilage in an aching joint currently remains unknown. Atypical innervation of joint tissues including the typically aneural cartilage has been reported in OA^6, 7^, and some studies suggested a correlation between sensory nerve fibre innervation, OA severity and OA-related pain^7-9^. These nerve fibres secrete sensory neuropeptides, such as α *calcitonin gene related peptide* (αCGRP), a 37 amino acid peptide best known as pain-mediator^10^. Of note, previous studies have reported elevated levels of αCGRP in serum, plasma and synovial fluid from patients with painful knee OA^11-14^, indicating that there are several potential cellular sources of αCGRP within the (OA) joint. Whilst αCGRP levels were suggested to correlate with OA pain and severity, sensory neuropeptides including αCGRP have also been described to act as trophic factors for several cell types of the joint, including osteoblasts and chondrocytes^15-17^, lending support to a potential active role of αCGRP as a driver of OA pathology, and a decrease in load resilience.

Some insights into potential functions of αCGRP in joint homeostasis and pathology come from recent animal and in vitro studies. Mice deficient for αCGRP were protected from age-related knee OA^18^, whereas cartilage degeneration was accelerated following surgical induction of OA in the absence of αCGRP^17^. Nevertheless, pharmacological inhibition of αCGRP via receptor antagonists attenuated OA progression following surgical OA induction^19, 20^, suggesting a context-dependent function of αCGRP in cartilage. When surgical OA-induction was followed by intense exercise, extracellular matrix (ECM) stiffening was prevented in αCGRP-deficient mice^21^, revealing effects of αCGRP on ECM organisation and biomechanical properties in vivo. Previous work reported a link between the mechano-response of cartilage and the sensory neuropeptide Substance P^22^, and so we asked whether the molecular response to mechanical loading may be altered by the presence of αCGRP.

Studies on the chondrocyte loading response typically employ cartilage explants or tissue-engineered 3D cartilage subjected to a strictly defined loading protocol in vitro. We have previously established a loading regimen which elicits an anabolic response in tissue-engineered cartilage, as demonstrated by increased proteoglycan synthesis rates on the metabolic level and by whole-transcriptome analysis^23^. In addition to increased extracellular signal-regulated kinase (ERK) phosphorylation and elevated expression of *FOS* and *PTGS2*, which are mechano-regulated in various tissues^24-28^, we also demonstrated enhanced *BMP2* and *BMP6* expression, and increased SRY-box transcription factor 9 (SOX9) protein levels, in line with the observed anabolic response of chondrocytes to the chosen loading regimen^23^. In agreement with studies employing different models and loading protocols^29^, we also reported that the cartilage ECM is a crucial parameter of the chondrocyte loading response^30^. Importantly, tissue-engineered cartilage allows the use of human cells and the accumulation of cartilage-like ECM, whilst also permitting defined culture and mechanical loading conditions, therefore filling an important gap between highly complex animal models and classical 2D cell culture experiments.

Here, we used this established model system to address whether αCGRP alters the molecular loading response of chondrocytes, and whether this can contribute to a decreased resilience of cartilage towards mechanical loading by investigating mechano-regulated signalling and the metabolic response of human chondrocytes in the presence of αCGRP. Tissue engineered neocartilage generated from human articular chondrocytes was exposed to αCGRP and subjected to a physiologically relevant anabolic loading regimen known to stimulate glycosaminoglycan (GAG) synthesis^23, 30^. The molecular response of untreated or αCGRP-treated neocartilage was then examined by qRT-PCR, western blotting and radiolabelling. A direct functional effect of αCGRP on the chondrocyte response to physiologically relevant mechanical stimulation has the potential to lead to novel approaches for OA treatment.

## METHODS

### Isolation and expansion of articular chondrocytes

Articular cartilage samples were obtained from 14 donors (3 males, 11 females, 52 – 87 years of age, mean age 66.9 years) undergoing total knee replacement surgery with informed written consent of the patients. The study was approved by the local ethics committee on human experimentation of the Medical Faculty of Heidelberg and in agreement with the Helsinki Declaration from 1975 (latest version). Articular chondrocytes (ACs) were isolated from macroscopically intact cartilage as described previously^23, 30^. Cells were expanded in DMEM containing 1 g/L glucose, 10 % FBS, 1 % penicillin/streptomycin and 2 mM L-glutamine (all Gibco^TM^, ThermoFisher Scientific, Germany) for two passages.

### Generation of tissue-engineered neocartilage

In order to generate neocartilage, ACs were seeded onto type I/III collagen carriers (4 mm diameter, 1.5 mm height, Optimaix, Matricel, Germany) at 5×10^5^ cells/construct, attached to a glass carrier (ROBU, Germany) to mimic the subchondral bone (bone replacement phase) and cultured for 35 days as described previously^23, 30^ using chondrogenic medium: DMEM (4.5 g/L glucose, 2 mM L-glutamine, Gibco^TM^, ThermoFisher Scientific), 1 % penicillin/streptomycin (Gibco^TM^, ThermoFisher Scientific), 0.1 mM dexamethasone, 0.17 mM ascorbic acid-2-phosphate, 2 mM sodium pyruvate, 0.35 mM proline, 5 µg/mL transferrin, 5 ng/mL sodium selenite, 1.25 mg/mL bovine serum albumin (all Sigma-Aldrich, Germany), 5 µg/mL insulin (Lantus, Sanofi-Aventis, Germany) and 10 ng/mL TGF-β1 (#100-21-100UG, Peprotech, Germany). 1 µM αCGRP (#4013281, Bachem, Germany) was added for the last 24 h of culture where indicated.

### Cyclic unconfined dynamic compression of neocartilage

Neocartilage was exposed to a single 3 h episode of mechanical loading in a custom-designed bioreactor with the dynamic compression protocol carried out using the software Galil Tools as described previously^23, 30^. In brief, neocartilage was initially compressed by 10 % of its thickness to ensure the constant contact with the piston during compression. Starting from this 10 % static offset, neocartilage was compressed by 25 % at 1 Hz during the loading intervals (10 min) whilst maintaining the offset of 10 % during break intervals (10 min). After loading, samples were washed in PBS and snap frozen, or used for metabolic labelling analysis. 1 µM αCGRP and where indicated 2 µM IWP-2 (Tocris, Germany) were added 21 h before start of compression and treatment was continued during loading and subsequent metabolic labelling.

### Radiolabel incorporation, glycosaminoglycan and DNA quantification

GAG synthesis was analysed after the end of loading as described previously^30^. Engineered tissue was labelled with 4 µCi ^35^S as Na_2_SO_4_ (Hartmann Analytic, Germany) for 24 h in 500 µL chondrogenic medium with or without treatments. Labelled neocartilage was then washed and digested with proteinase K (ThermoFisher Scientific) overnight. Incorporated radiolabel was measured in tissue lysates by β-scintillation counting (1414 WinSpectral, PerkinElmer, Germany) and normalised to the DNA content of the same lysates. DNA content was determined using Quant iT Pico Green Kit (Invitrogen, Germany). Total GAG content of each sample was determined by 1,9-dimethylmethylene blue (DMMB) assay using a chondroitin-6-sulfate standard (Sigma-Aldrich).

### RNA isolation and mRNA expression analysis

Total RNA was isolated from neocartilage using Trizol (Life Technologies, Germany). 500 ng of total RNA was reverse transcribed using Omniscript RT Kit (Qiagen, Germany) and oligo(dT) primers. Gene expression was analysed by qRT-PCR (Roche Diagnostics, Germany) using SYBR green (ThermoFisher Scientific). *CPSF6* and *HNRPH1* were used as housekeeping genes. Primer sequences are listed in table S1.

### Western blotting

Neocartilage was cut in half and ground in PhosphoSafe Extraction Reagent (Merck Millipore, Germany) containing 1 mM Pefabloc SC (Sigma-Aldrich, Germany) in a mixer mill (Retsch, Germany) at 30 Hz for 2 x 2 min, with cooling on ice in between cycles. Samples were centrifuged at 13000 x g for 20 min at 4°C before cell lysates were mixed with 4 x Laemmli buffer and boiled for 5 min at 95°C. Proteins were separated according to size using standard SDS-PAGE and transferred onto a nitrocellulose membrane (GE Healthcare, Amersham, Germany). Membranes were blocked in 5 % skimmed milk/TBS-T and probed with primary antibodies (see table S2) at 4°C overnight. Detection was carried out using horseradish peroxidase-conjugated secondary antibodies (see table S2) and enhanced chemiluminescence (Roche Diagnostics, Germany/Advansta, USA). Densitometric analysis was performed using ImageJ.

### Histology

Neocartilage was separated from the bone replacement phase and fixed in 4% PFA for 4 h at room temperature. After dehydration in an ascending isopropanol series, samples were embedded in paraffin wax for slide preparation. After rehydration, proteoglycan deposition was analysed by staining 5 µm sections with 0.2 % safranin-O (Fluka, Sigma-Aldrich, Germany) in 1 % acetic acid and counterstaining with 0.04 % Certistain fast green (Merck, Germany) in 0.2 % acetic acid. Type II collagen immunostaining was performed as follows: Sections were rehydrated and digested with hyaluronidase (Roche Diagnostics, Mannheim, Germany, 4 mg/mL in PBS, pH 5.5) followed by pronase (Roche Diagnostics, 1 mg/mL in PBS, pH 7.4), before unspecific binding sites were blocked using 5 % BSA. After incubation with mouse anti-human type II collagen antibody (see table S2), sections were incubated with ALP-coupled anti-mouse secondary antibody (see table S2), and ImmPACT^®^ Vector^®^ Red Substrate (Vector Laboratories, Newark, CA, USA). As negative controls, undifferentiated stem cells cultured in a collagen carrier were used, in addition to a type II collagen-positive neocartilage sample stained without primary antibody (see Fig. S1B).

### Statistical analysis

Statistical analysis was performed using IBM SPSS Statistics (29.0.0.0). Fold changes were calculated relative to the respective free-swelling samples and *P*-values calculated using appropriate statistical testing as noted in the figure legends, with *P* < 0.05 considered statistically significant.

## RESULTS

### Neocartilage quality is sustained in the presence of αCGRP

In order to investigate the effect of αCGRP on the immediate molecular response of chondrocytes to mechanical loading, we chose to limit the αCGRP treatment to the last 24 h of culture, including the dynamic unconfined compression on day 35, thus avoiding previously reported reductions in GAG deposition following αCGRP treatment for 7 days^31^. At the end of culture, untreated and αCGRP-treated neocartilage was histologically indistinguishable (Fig. 1A), with both groups exhibiting comparable, strong GAG and type II collagen staining. Accordingly, the GAG synthesis rate as well as total GAG content was generally maintained in the presence of αCGRP (Fig. 1B). No significant changes were observed in the DNA content of control or αCGRP-treated neocartilage (Fig. S1A). Similar expression levels of the chondrocyte marker genes *COL2A1*, *ACAN* and *SOX9* between control and αCGRP-treated samples (Fig. 1C) also indicated that short-term αCGRP treatment did not exert significant negative effects on neocartilage.

**Figure 1.**
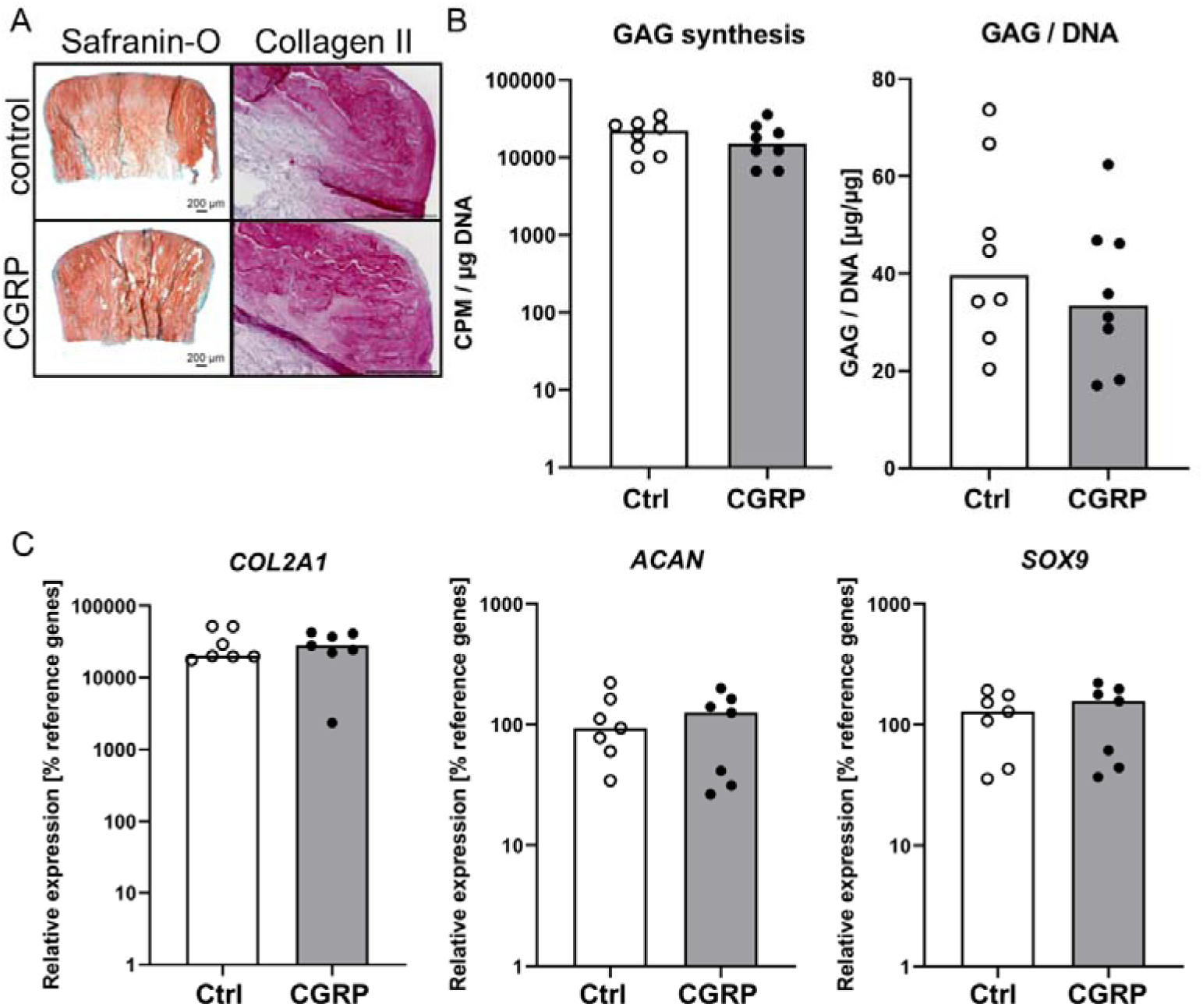
Neocartilage ECM structure and quality is sustained in the presence of αCGRP. Articular chondrocytes were cultured in collagen carriers for 35 days and treated with 1 µM αCGRP for the last 24 h of culture. (A) Histological staining of glycosaminoglycans (GAG) by safranin-O/fast green staining and of type II collagen by immunohistochemistry (IHC). Positive and negative controls for IHC are provided in the supplements (Fig. S1B). N = 5 donors. Scale bar: 500 µM unless otherwise stated. (B) GAG synthesis over 24 h was assessed by radiolabelling using ^35^S-sulfate (Na_2_^35^SO_4_). Changes in ^35^S-sulfate incorporation were normalised to DNA content (measured by PicoGreen assay). GAG content was determined using the DMMB method and normalised to the DNA content. N = 8 donors. (C) Expression of chondrocyte marker genes *COL2A1, ACAN,* and *SOX9* was quantified by qRT-PCR using *CPSF6* and *HNRPH1* as reference genes. N = 7 donors. Bars represent the median value.

### αCGRP does not impair chondrocyte mechano-transduction

Next, day 35 neocartilage tissue constructs were subjected to a previously established anabolic loading protocol consisting of 3 h of dynamic unconfined compression in the presence or absence of αCGRP. In order to assess the chondrocyte mechano-transduction response, phosphorylation of the MAP kinase ERK1/2 was examined by western blotting (Fig. 2A). As expected, mechanical loading significantly enhanced ERK1/2 phosphorylation in the control group. ERK1/2 phosphorylation was also significantly stimulated in the presence of αCGRP. We then examined expression levels of known mechano-regulated genes, including *FOS* and *PTGS2* (Fig. 2B). Expression of both genes was significantly enhanced following mechanical loading, in the control group (median fold change 4.61 and 4.72, respectively) as well as in the presence of αCGRP (median fold change 4.12 and 3.91, respectively). Furthermore, SOX9 protein levels were significantly increased by mechanical loading, both in the presence as well as absence of αCGRP (Fig. 2C). This was also reflected in upregulated mRNA levels (Fig. S2), in line with our previous reports^23^. Furthermore, *BMP2* and *BMP6* expression was stimulated by mechanical loading in both groups (Fig. 2D). Since these known mechano-regulated factors still responded to compression in the presence of αCGRP, we concluded that chondrocyte mechano-transduction was generally not affected by the presence of αCGRP.

**Figure 2.**
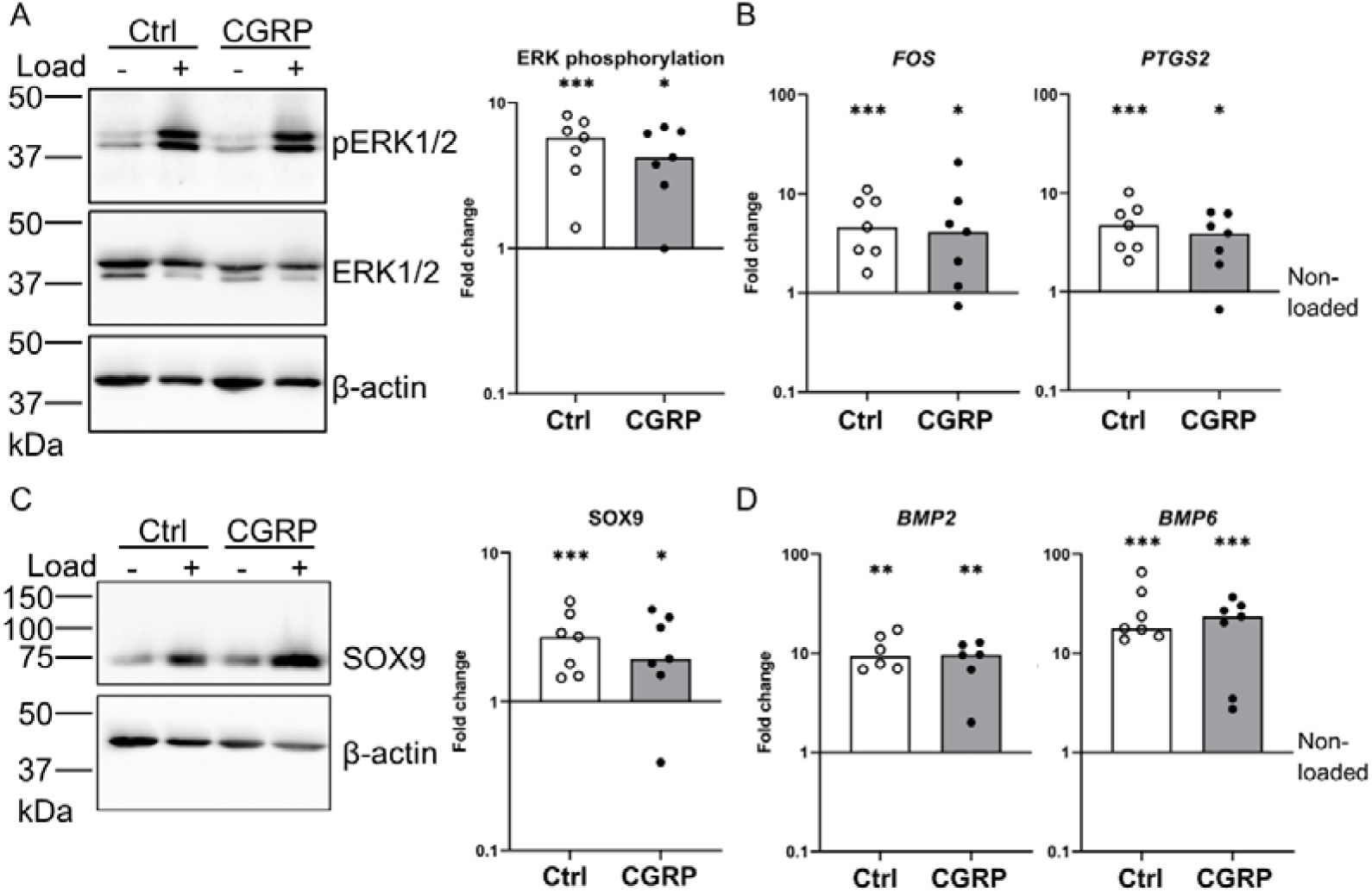
Chondrocyte mechano-transduction is not altered by αCGRP. Neocartilage was subjected to 3 h of mechanical loading in the presence or absence of 1 µM αCGRP, added for the last 24 h of culture (including the loading period). (A) Visualisation and quantification of ERK phosphorylation by western blotting. Densitometric analysis was performed in ImageJ using β-actin as loading control and the fold change calculated relative to the non-loaded control set to 1. (B) Gene expression analysis of *FOS* and *PTGS2* by qRT-PCR, using *CPSF6* and *HNRPH1* as reference genes, and fold changes calculated relative to the non-loaded control set to 1. (C) Visualisation and quantification of SOX9 protein levels by western blotting. Densitometric analysis was performed as above. (D) Gene expression analysis of *BMP2* and *BMP6* by qRT-PCR, using *CPSF6* and *HNRPH1* as reference genes, and fold changes calculated relative to the non-loaded control set to 1. N = 6-7 donors. Bars represent the median value. P-values were calculated using Mann-Whitney-U testing with the non-loaded control set to 1, * indicate p < 0.05, ** indicate p < 0.01, *** indicate p < 0.001. Uncropped images of western blot membranes are provided in the supplemental information (Fig. S5).

### Load-stimulated proteoglycan production is disturbed by αCGRP

The impact of αCGRP treatment on the functional response of chondrocytes was assessed by performing ^35^S-sulfate isotope labelling of newly synthesised GAGs over 24 h after the end of loading (Fig. 3A). In the control group, dynamic compression resulted in significantly enhanced GAG synthesis by an average of 24.7 %, a value in agreement with our previous studies^30^. Although no major alterations in mechano-transduction or mechano-regulated gene expression were observed after αCGRP treatment, the stimulation of GAG synthesis following loading was clearly disrupted (Fig. 3A). DNA content per sample and total GAG content were maintained in both control and αCGRP-treated neocartilage following loading, indicating that the αCGRP effect was not due to tissue disruption (Fig. 3B, C). Furthermore, the effect of αCGRP on load-mediated GAG synthesis stimulation appeared dose-dependent in a smaller set of experiments (Fig. S3).

**Figure 3.**
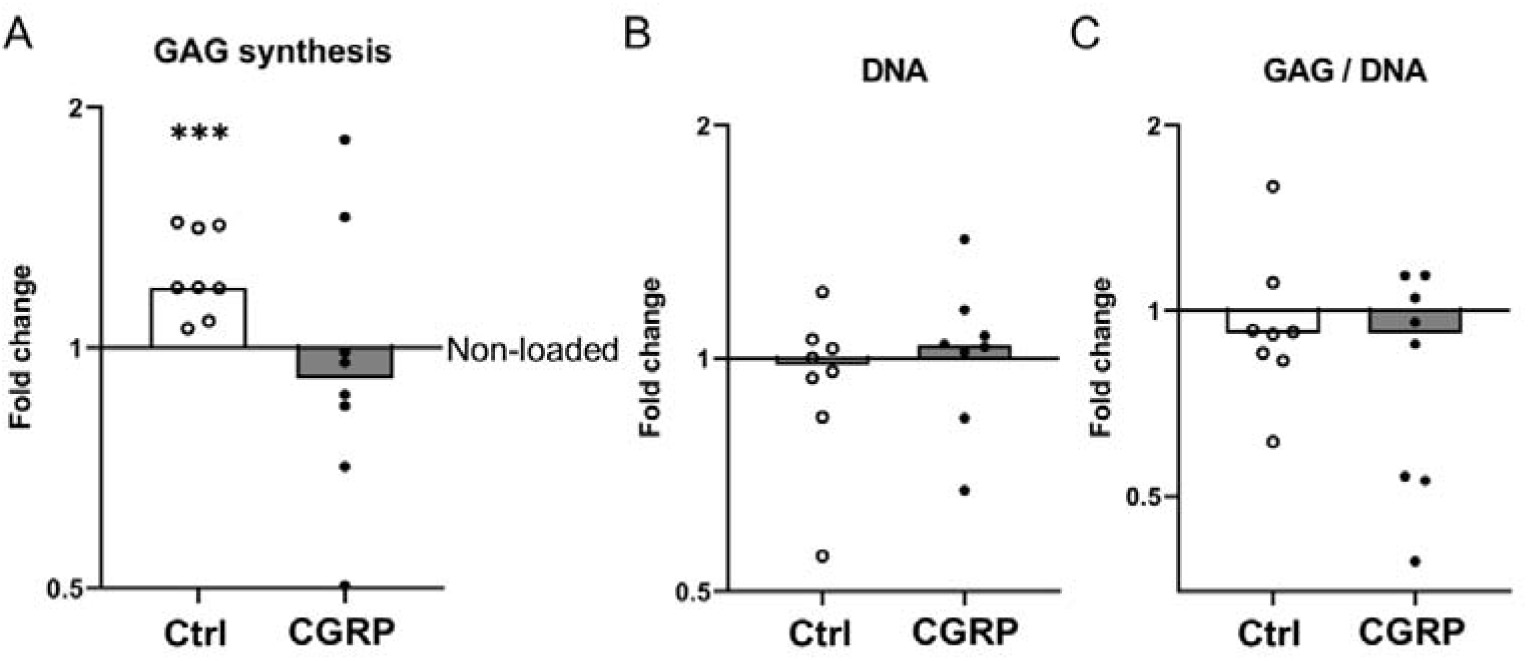
Load-stimulated GAG synthesis is disturbed by the presence of αCGRP. Neocartilage was subjected to 3 h of mechanical loading in the presence or absence of 1 µM αCGRP, added for the last 24 h of culture. (A) GAG synthesis was assessed by radiolabelling for 24 h after loading using ^35^S-sulfate (Na_2_^35^SO_4_). ^35^S-sulfate incorporation was normalised to DNA content and referred to the non-loaded control set to 1. (B) DNA content, measured by PicoGreen assay was assessed following radiolabelling and normalised to the non-loaded control set to 1. (C). Total GAG content was quantified by the DMMB method, normalised to the DNA content per sample, and referred to the non-loaded control set to 1. N = 8 donors. Bars represent the median value, p-values were calculated using Mann-Whitney-U testing with the control set to 1, *** indicate p < 0.001.

### αCGRP-induced disruption of proteoglycan production after mechanical loading is alleviated by WNT inhibition

We then explored the mechanism by which αCGRP interferes with load-induced GAG synthesis. Our previous studies identified a precise regulation of WNT activity as crucial factor for the chondrocyte loading response^30^ and in human osteoblasts, treatment with αCGRP was previously described to result in the stabilisation of the central WNT mediator β-catenin^16^. Therefore, we next examined whether the presence of αCGRP during mechanical loading affects the WNT signalling network of human neocartilage. Amongst WNT ligands expressed in human chondrocytes according to our previously published data^32^, WNT5A inhibited anabolic gene expression in human chondrocytes^33^. Interestingly, several studies on human tenocytes revealed an upregulation of *WNT5A* in response to mechanical stimulation^34, 35^. Thus, we examined whether gene expression of *WNT5A* was regulated following mechanical loading in control and αCGRP-treated neocartilage (Fig. 4A). Indeed, *WNT5A* mRNA levels were significantly elevated following mechanical loading in both groups (median fold change 1.38 and 1.42, respectively). Of note, the amount of another chondrocyte-expressed WNT ligand (*WNT11*)^32^ remained unchanged by mechanical loading in the presence or absence of αCGRP. We also examined the expression of the WNT antagonist *DKK3,* the only expressed form of dickkopf under the chosen conditions according to our previous array data^32^. DKK3 has previously been described to prevent cytokine-induced proteoglycan loss and WNT-induced proteoglycan reduction^36^. Whilst mechanical loading had no significant effect on *DKK3* expression in the non-treated control group (median fold change 0.86), *DKK3* transcript levels were significantly reduced by nearly 40 % when mechanical loading was performed in the presence of αCGRP (median fold change 0.61, p < 0.001). Treatment with αCGRP alone did not significantly alter expression of *WNT5A, WNT11* or *DKK3* compared to non-loaded control neocartilage (Fig. S4A).

**Figure 4.**
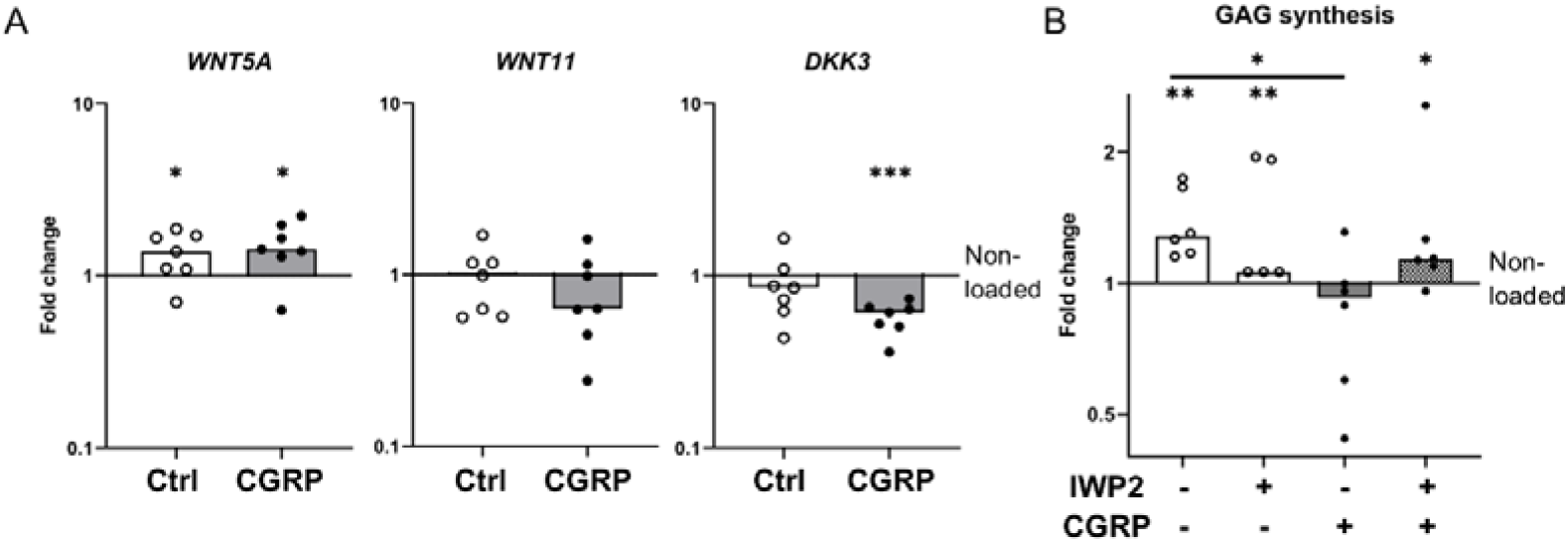
WNT inhibition alleviates αCGRP-mediated restriction of proteoglycan production. Neocartilage was subjected to 3 h of mechanical loading with 1 µM αCGRP and 2 µM IWP-2 added for the last 24 h of culture where indicated. (A) Gene expression analysis by qRT-PCR, using *CPSF6* and *HNRPH1* as reference genes and fold changes calculated relative to the non-loaded control set to 1. N = 7 donors. (B) GAG synthesis was assessed by radiolabelling using ^35^S-sulfate (Na_2_^35^SO_4_). Changes in ^35^S-sulfate incorporation were normalised to DNA content and referred to the non-loaded control set to 1. Bars represent the median value. N = 5-6 from 4-5 donors. P-values were calculated using Mann-Whitney-U testing with the control set to 1, * indicate p < 0.05, ** indicate p < 0.01, *** indicate p < 0.001.

In order to investigate whether αCGRP exerts its effect on proteoglycan production via modulation of WNT signalling, neocartilage was subjected to dynamic compression in the presence of αCGRP, and additionally treated with the pan-WNT inhibitor IWP-2 (2 µM) or DMSO as solvent control. Neither DNA content nor GAG/DNA ratio were significantly altered following IWP-2 treatment, αCGRP treatment, or co-treatment (Fig. S4B, C). As expected, GAG synthesis was stimulated by mechanical loading in the control group (on average by 38%), as well as in the presence of IWP-2, whereas αCGRP prevented this response (Fig. 4B). Strikingly, the inhibitory effect of αCGRP on the stimulation of GAG synthesis following compression was alleviated in the presence of IWP-2 (average stimulation of 36 %). Together, these data reveal that αCGRP can disturb chondrocyte proteoglycan production in response to anabolic loading by interfering with the delicate balance of WNT signalling activity in cartilage. Thus αCGRP at least partly impairs the mechano-adaptive response of chondrocytes to mechanical stimulation.

## DISCUSSION

Responding adequately to mechanical loading is an important function of articular cartilage, which is impaired by degenerative processes during OA and thus renders cartilage less resilient towards mechanical challenge. Illuminating the effects of OA-relevant changes on the response of chondrocytes to mechanical stimulation has the potential to identify novel causative therapeutic targets and may suggest new strategies of stratifying patients who can especially benefit from a specific therapy. Previous reports suggested a correlation between αCGRP levels and OA pain^11, 13, 14^, and in vivo studies employing mouse models revealed roles for αCGRP also in structural aspects of joint pathology^15, 17, 18, 21^. Yet, the effects of αCGRP on functional aspects of cartilage, such as the chondrocyte loading response, have remained so far unaddressed. In our present study, we reveal that αCGRP can compromise the anabolic response of chondrocytes to a physiological loading regimen, and thus the resilience of cartilage towards mechanical loading. We further show that WNT inhibition alleviates this effect of αCGRP, altogether suggesting αCGRP as a potential target for OA treatment.

Our study employed a previously established and well characterised tissue-engineering model based on human chondrocytes^23, 30^, which allows their mechanical stimulation under defined experimental conditions. Thus, we were able to investigate important functional aspects of human cartilage biology, thereby filling the large gap between the complex in vivo environment and oversimplified (2D) cell culture models. By limiting the duration of αCGRP exposure, we prevented significant effects on proteoglycan accumulation by αCGRP treatment alone as described by an earlier study^31^, allowing us to identify how αCGRP influences the chondrocyte loading response. Importantly, the selected loading regimen reflects physiological loading conditions by employing an intermittent compression protocol, and presents an anabolic stimulus^23^. This model therefore provided the exciting opportunity to assess whether αCGRP can alter the metabolic response of chondrocytes to physiological mechanical loading under defined and strictly-controlled conditions.

Using this model, we demonstrated for the first time that αCGRP disrupted the stimulation of GAG synthesis by mechanical loading, suppressing the anabolic effect of this physiological loading regimen. Our current data provide novel and compelling evidence to implicate αCGRP in a reduced load resilience of cartilage, further supporting functional contributions of sensory neuropeptides to cartilage homeostasis, but also to cartilage degeneration during OA and suggesting αCGRP as a potential therapeutic target. Specific αCGRP-targeted therapeutics are already approved for unrelated conditions including migraines (as reviewed in^37^). Our findings are in line with previous studies using human chondrocytes under static conditions, which revealed reduced GAG deposition in the presence of αCGRP^31^ as well as studies employing animal models, which linked αCGRP to changes in ECM organisation and biomechanical properties^17, 21^. In vivo studies also revealed that αCGRP-deficiency protected knee cartilage from degradation in a mouse model of age-related OA^18^, and pharmacological inhibition of αCGRP also attenuated cartilage degeneration following surgical induction of OA^19, 20^. Yet, life-long αCGRP-deficiency enhanced cartilage degeneration following induction of post-traumatic OA^17^, and intra-articular injection of αCGRP-overexpressing bone marrow stromal cells following surgical OA induction appeared to have beneficial effects on general mobility and activity in a mouse model^38^. Genetic ablation of αCGRP is known to result in osteopenia^15^, and altered homeostatic conditions within the joint may explain these discrepancies. Still, further studies into the context-dependent regulation of αCGRP as an active driver of OA pathology are required to resolve these apparent contradictions.

Importantly, our study linked the effect of αCGRP in the context of mechanical loading to WNT activity. The WNT signalling pathway is well-known for its contribution to cartilage degeneration and OA pathology^39^ and during our previous study, stabilisation of the WNT-mediator β-catenin prevented the anabolic response of chondrocytes to the chosen loading episode^30^. Conversely, a pan-WNT inhibitor rescued the load-induced reduction of GAG synthesis in presence of high WNT/β-catenin levels as a potent negative regulator of load-induced chondrocyte anabolism. Additionally, αCGRP was previously reported to contribute to a stabilisation of β-catenin in osteoblasts^16^, in line with pro-anabolic effects of αCGRP in bone in vivo^15^. Here, we applied this knowledge to mitigate the effect of αCGRP on GAG synthesis stimulation following mechanical loading, using the WNT inhibitor IWP-2. Since IWP-2 acts via blocking the secretion of endogenously produced WNT ligands, our data implicate a WNT ligand-mediated pathway in the action of αCGRP.

Which WNT ligands are involved in chondrocyte mechano-transduction and in mediating the effect of αCGRP remains to be elucidated in future studies. Here, we report for the first time elevated expression of the non-canonical WNT ligand *WNT5A* as a novel mechano-response gene of human chondrocytes. *WNT5A* was recently identified as an effector gene in osteoarthritis^40^. Increased expression of *WNT5A* has also been described in response to cyclic stretch and tensile strain in human tenocytes as well as human mesenchymal stromal cell-derived tenocytes and murine superficial zone chondrocytes^34, 41^. Importantly, our previous transcriptome-wide study^23^ revealed the mechano-regulation of known WNT target genes, but the responsible ligand remains unknown. Other studies have also described WNT/β-catenin activity following mechanical loading of chondrocytes^42^; however, the mediating ligand was also not identified. It remains to be addressed by future studies whether the mechano-regulation of *WNT5A* contributes to WNT activity in chondrocytes, and whether this involves β-catenin-dependent or -independent mechanisms. Importantly, in αCGRP-treated neocartilage, mechanical loading resulted in a concurrent decrease in *DKK3* expression. Depending on the biological context, DKK3 has been described as non-canonical or canonical WNT inhibitor; however, at least one previous study has shown that DKK3 can antagonise the archetypal canonical WNT ligand WNT3A in human chondrocytes^36^. It is tempting to speculate that a reduction in *DKK3* expression, but sustained upregulation of *WNT5A* in the presence of αCGRP could result in an altered balance of non-canonical vs canonical WNT activity following mechanical loading, and that this could contribute to a decreased resilience of cartilage towards mechanical loading.

Our findings implicating WNT activity as a mediator of αCGRP could also be relevant for patient stratification. Whilst patient heterogeneity as well as insufficient stratification of patient subgroups and phenotypes remain a major problem in the development and clinical testing of DMOADs, one of the very few substances that have progressed into advanced clinical trials (reviewed in^43^), lorecivivint, was described to modulate WNT activity^44, 45^. Thus, it would be interesting to investigate whether lorecivivint is able to mitigate the effect of αCGRP on proteoglycan production following mechanical loading. This knowledge could link the potential DMOAD to an improved resilience of cartilage towards mechanical loading. Since αCGRP has already successfully been detected in synovial fluid and serum samples from OA patients, increased αCGRP levels could possibly identify patients that would benefit particularly from WNT activity modulation.

Our study has several limitations. Since we performed the αCGRP treatment after formation of a cartilage ECM, we used a supra-physiological concentration of αCGRP, which is higher than concentrations used in other studies using human chondrocytes^31^. However, in contrast to other studies, which have applied αCGRP during the entire culture period, we added αCGRP to already matured neocartilage and therefore raised the concentration in order to ensure penetration of the dense, collagen- and proteoglycan-rich extracellular matrix. Another limitation is that we tested the effect of αCGRP treatment only in combination with one physiological loading regime. Future studies are required to assess the effects of other types of loading protocols (such as mechanical overloading), and multi-axial loading conditions in the presence of αCGRP. It is also important to note that WNT activity-related β-catenin is typically analysed following the saponin extraction method^46^ to eliminate membrane-bound β-catenin involved in cell adhesion; however, the dense structure of (tissue-engineered) cartilage is incompatible with this extraction method, which precluded the analysis of β-catenin levels. Lastly, we relied on the generous tissue donations from OA patients undergoing total knee replacement surgery, so chondrocytes were derived from OA patients; however, we took this into account during experimental design and strictly limited exposure to αCGRP in order to prevent the effects on proteoglycan accumulation described for OA but not non-OA chondrocytes^31^.

In conclusion, we showed here for the first time that the sensory neuropeptide αCGRP, which was previously implicated in OA progression, can impair load-stimulated proteoglycan production by human chondrocytes, and thus interferes with their anabolic response to physiological mechanical loading. Our findings indicated that αCGRP can compromise the resilience of cartilage towards mechanical loading, and linked this effect of αCGRP to WNT activity. We also report *WNT5A* as a novel mechano-response gene in human chondrocytes. Overall, our data support a role for αCGRP in cartilage degeneration and OA progression and as a potential new therapeutic target. Elevated αCGRP levels may also help stratifying patients that can benefit from a WNT antagonistic treatment.

## Supporting information

Supplementary table 1 - primer sequences

Supplementary table 2 - antibodies

## Acknowledgements

We thank Dr. Christian Kleist and Nadine Weissheimer for their generous assistance with liquid scintillation counting, Barbara König, Elena Tripel and Sarina Losch for excellent technical assistance, the surgical staff for support with sample collection, and I. Heschel from Matricel for providing the Optimaix scaffolds. We would also like to thank Dr. Sabine Stöckl for helpful advice. This study was supported by the German Research Foundation DFG-grant R1707/12-2 and GR1301/19-2 as part of the Excarbon Research Group FOR2407, the University of Regensburg and the Orthopaedic University Hospital Heidelberg.

## CRediT authorship statement

**Helen F. Dietmar**: Investigation, Conceptualisation, Formal analysis, Data curation, Visualisation, Writing – original draft, Writing – review & editing. **Nicole Hecht**: Investigation, Conceptualisation, Methodology, Formal analysis, Data curation, Writing – review & editing. **Carina Binder**: Investigation, Methodology, Data curation, Writing – review & editing. **Sven Schmidt:** Formal analysis, Writing – review & editing. **Tilman Walker**: Resources, Funding acquisition, Writing – review & editing. **Susanne Grässel:** Conceptualisation, Project administration, Funding Acquisition, Resources, Writing – review & editing. **Wiltrud Richter**: Conceptualisation, Project administration, Supervision, Funding Acquisition, Resources, Writing – review & editing. **Solvig Diederichs**: Conceptualisation, Formal analysis, Project administration, Supervision, Funding Acquisition, Resources, Writing – original draft, Writing – review & editing.

