## Supplementary figures for "The neuropeptide alpha calcitonin gene-related peptide impairs load-stimulated proteoglycan production of human chondrocytes via WNT activity"

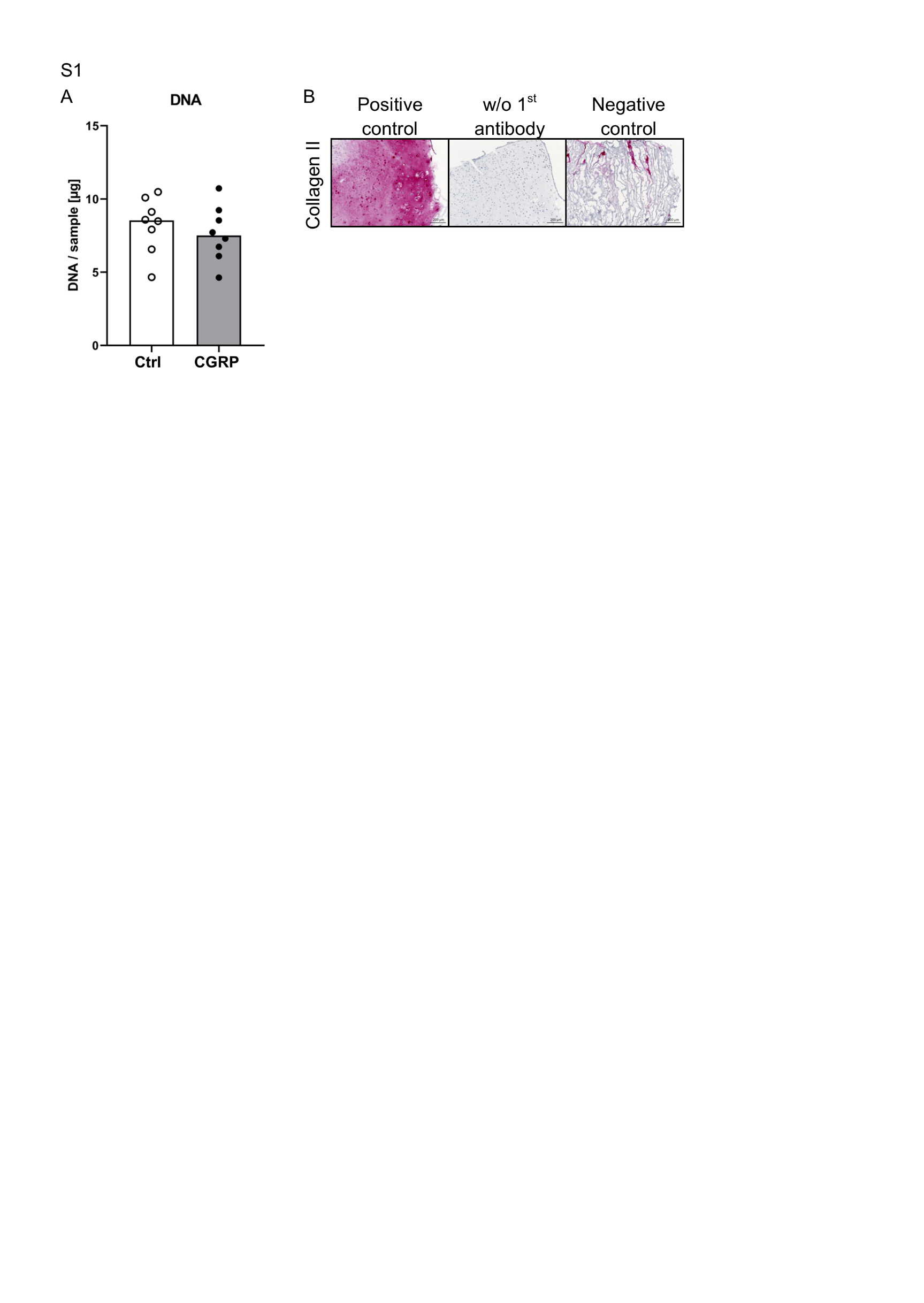

Figure S1

(A) Articular chondrocytes were cultured in collagen scaffolds for 35 days and treated with 1 µM αCGRP for the last 24 h of culture. DNA content as measured by PicoGreen assay and per neocartilage construct. N = 8 donors. Bars indicate median values. (B) Positive and negative controls for type II collagen IHC. Previously generated and analysed neocartilage served as positive control, and was also used for the secondary antibody-only (w/o 1^st^ antibody) control. Undifferentiated stem cells cultured in a collagen carrier served as negative control.

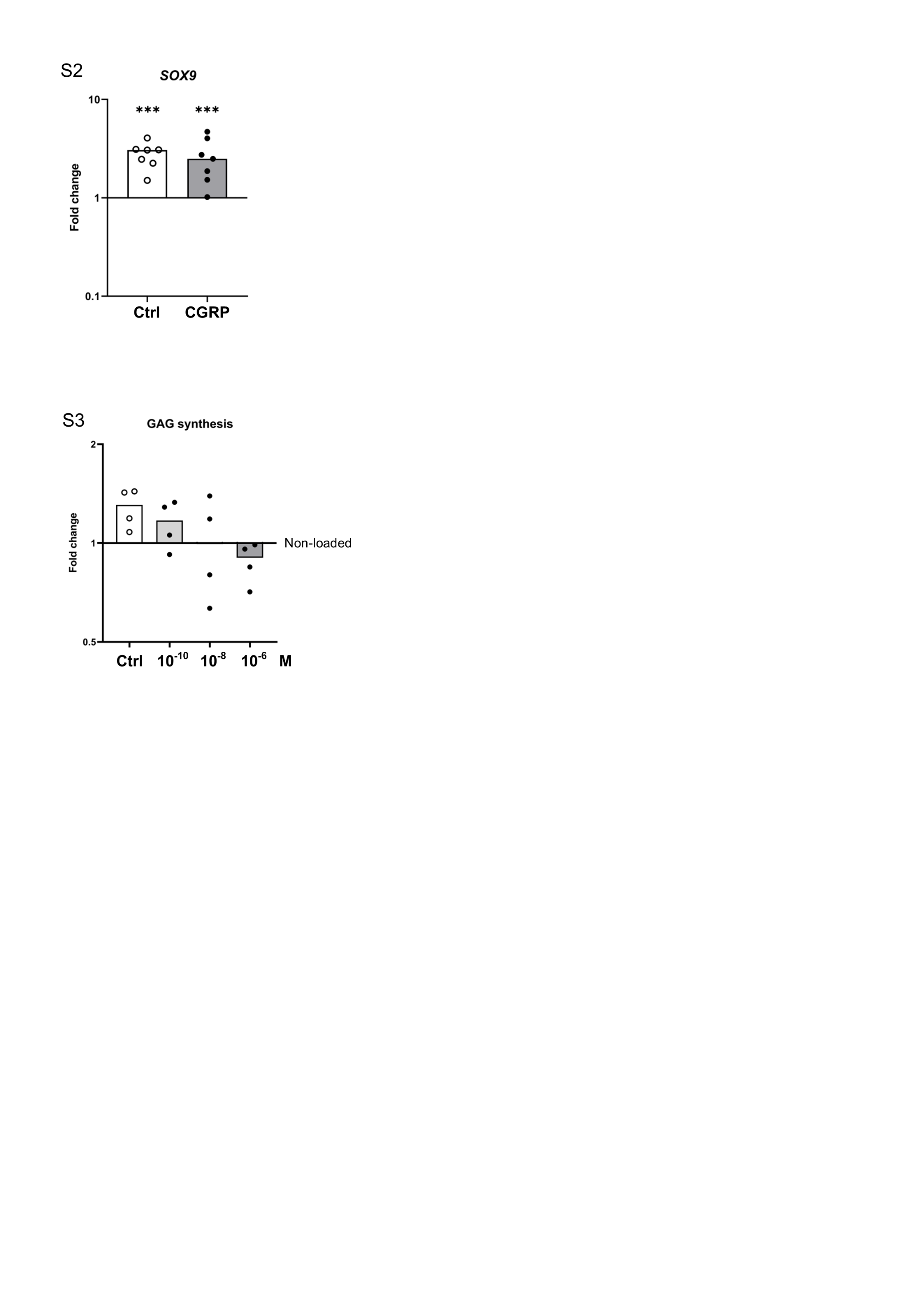

Figure S2 – Stimulation of *SOX9* mRNA is not altered by the presence of αCGRP during mechanical loading. Neocartilage was subjected to 3 h of mechanical loading in the presence or absence of 1 µM αCGRP, added for the last 24 h of culture. Gene expression of *SOX9* was analysed by qRT-PCR, using *CPSF6* and *HNRPH1* as reference genes, and fold changes calculated relative to the non-loaded control set to one. Bars indicate median values. N = 7 donors. P-values were calculated using Mann-Whitney-U testing with the control set to 1, *** indicate p < 0.001.

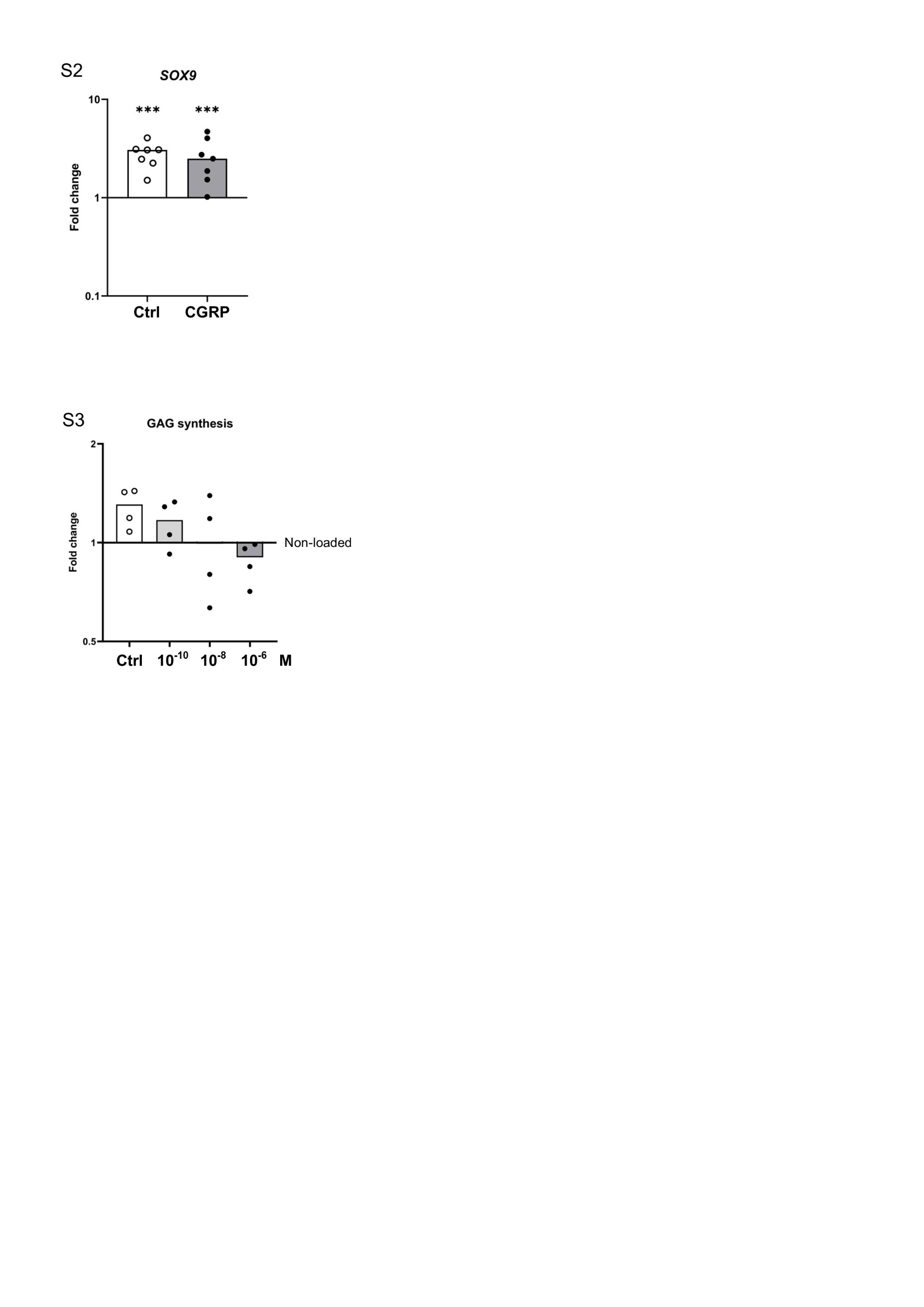

Figure S3 - Effects of αCGRP concentration on load-stimulated GAG synthesis rates. Neocartilage was subjected to 3 h of mechanical loading in the presence of the indicated concentration of αCGRP, added for the last 24 h of culture. GAG synthesis was assessed by radiolabelling using ^35^S-sulfate (Na_2_^35^SO_4_). Changes in ^35^S-sulfate incorporation were normalised to DNA content and referred to the non-loaded control set to 1. Bars indicate median values. N = 4 donors.

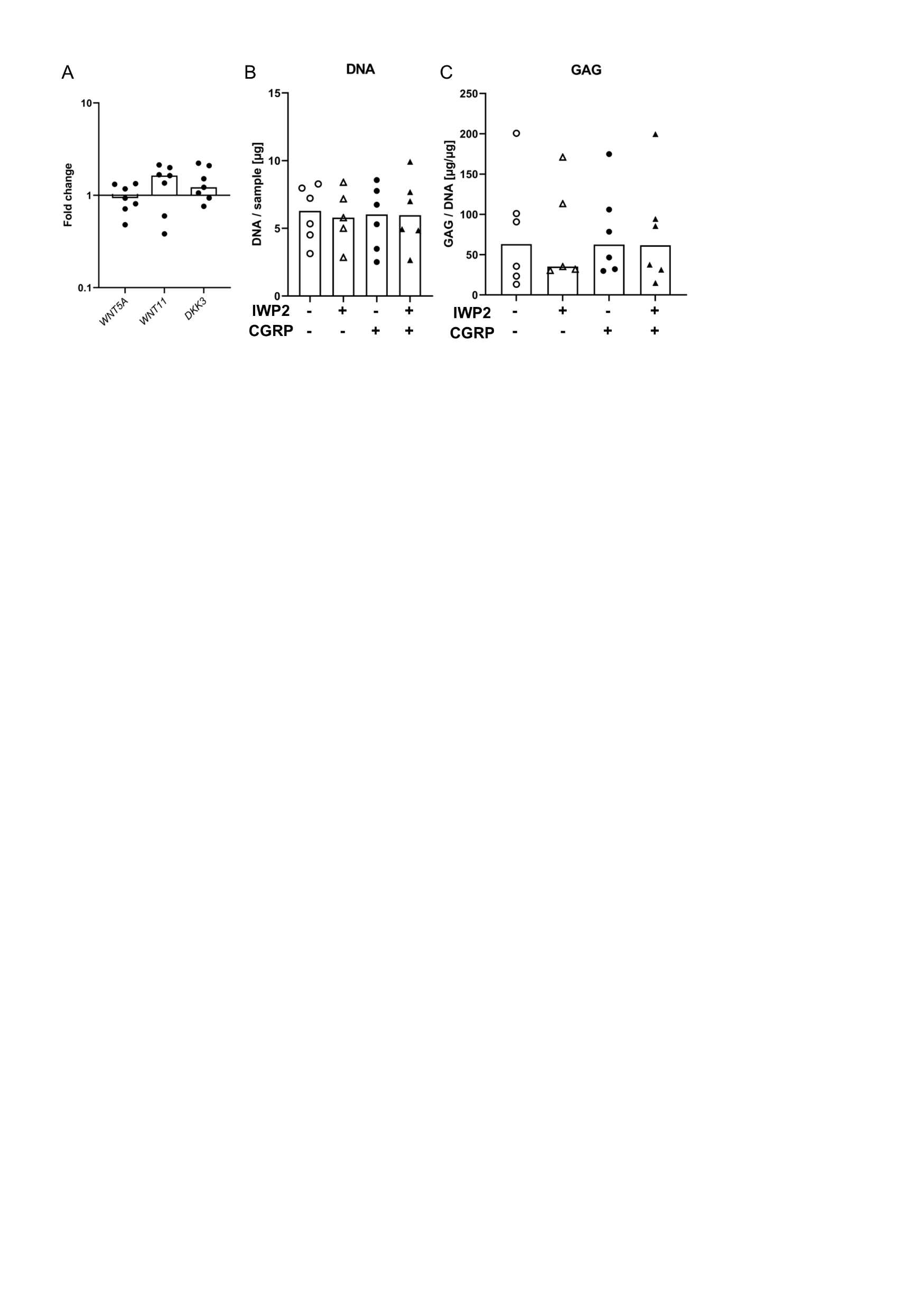

Figure S4

(A) Neocartilage was cultured for 35 days, with 1 µM αCGRP added for the last 24 h of culture. Gene expression of *WNT5A, WNT11* and *DKK3* was analysed by qRT-PCR, using *CPSF6* and *HNRPH1* as reference genes, and fold changes calculated relative to the untreated control set to 1. N = 7 donors. (B) DNA content, measured by PicoGreen assay was assessed following radiolabelling. N = 5-6 from 4-5 donors. (C). Total GAG content was quantified by DMMB method and normalised to the DNA content per sample. 1 µM αCGRP and/or 2 µM IWP-2 were added for the last 24 h of culture as indicated. N = 5-6 from 4-5 donors. Bars indicate median values.

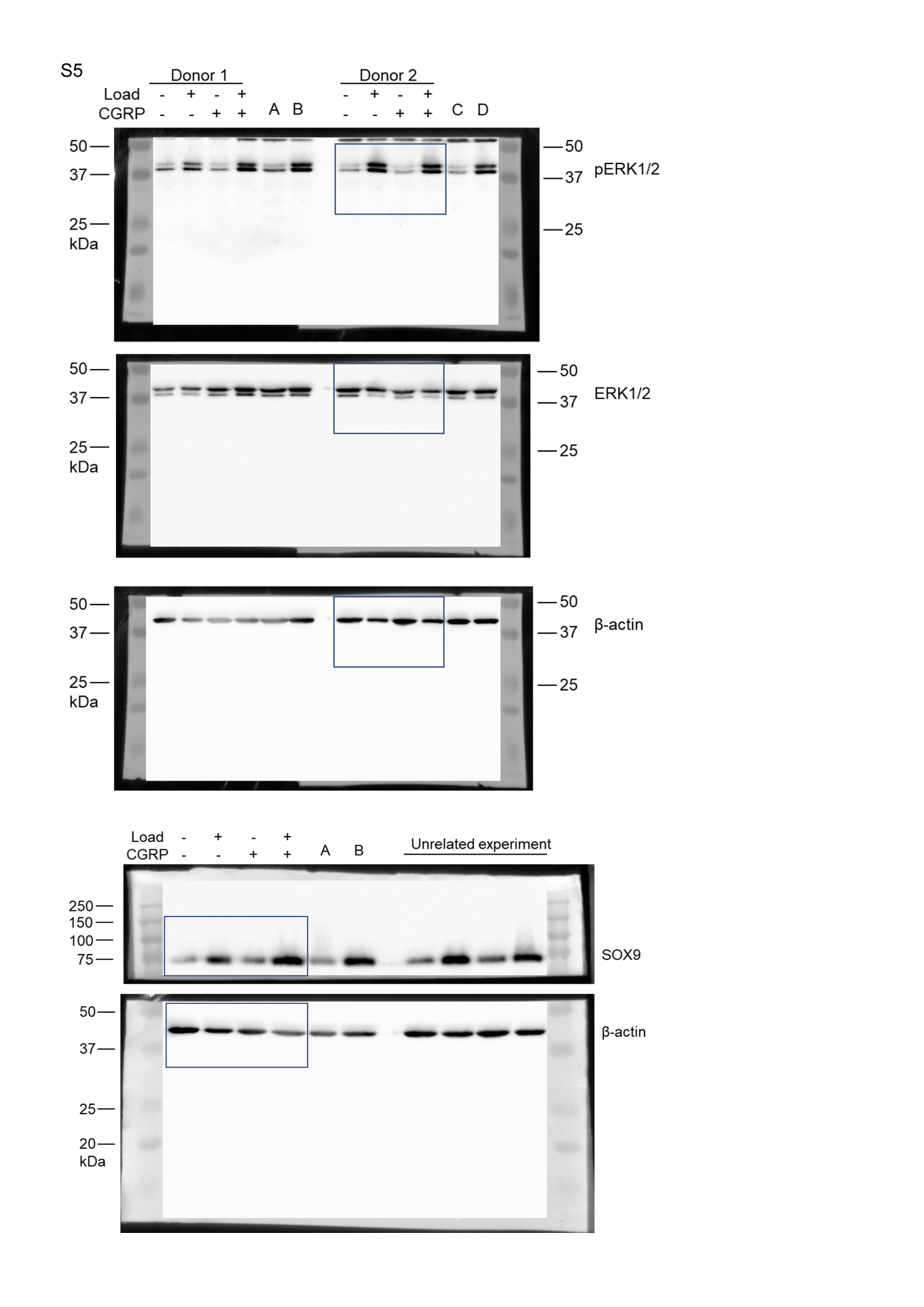

Figure S5 - Uncropped western blot membranes (shown in Fig. 2.).
